# An insight into SARS-CoV-2 Membrane protein interaction with Spike, Envelope, and Nucleocapsid proteins

**DOI:** 10.1101/2020.10.30.363002

**Authors:** Prateek Kumar, Amit Kumar, Neha Garg, Rajanish Giri

**Affiliations:** School of Basic Sciences, Indian Institute of Technology Mandi, VPO Kamand, Himachal Pradesh, 175005, India; Department of Medicinal Chemistry, Faculty of Ayurveda, Institute of Medical Sciences, Banaras Hindu University, Varanasi, Uttar Pradesh, 221005, India

**Keywords:** Protein-protein interactions, binding free energy, Molecular dynamics simulation, Membrane, Envelope, Nucleocapsid, Spike Glycoprotein

## Abstract

Intraviral protein-protein interactions are crucial for replication, pathogenicity, and viral assembly. Among these, virus assembly is a critical step as it regulates the arrangements of viral structural proteins and helps in the encapsulation of genomic material. SARS-CoV-2 structural proteins play an essential role in the self-rearrangement, RNA encapsulation, and mature virus particle formation. In SARS-CoV, the membrane protein interacts with the envelope and spike protein in Endoplasmic Reticulum Golgi Intermediate Complex (ERGIC) to form an assembly in the lipid bilayer, followed by membrane-ribonucleoprotein (nucleocapsid) interaction. In this study, we tried to understand the interaction of membrane protein’s interaction with envelope, spike, and nucleocapsid proteins using protein-protein docking. Further, simulation studies performed up to 100 ns to examine the stability of protein-protein complexes of Membrane-Envelope, Membrane-Spike, and Membrane-Nucleocapsid. Prime MM-GBSA showed high binding energy calculations than the docked complex. The interactions identified in our study will be of great importance, as it provides valuable insight into the protein-protein complex, which could be the potential drug targets for future studies.

## Introduction

Seven types of coronaviruses infect humans, among which severe acute respiratory syndrome (SARS-CoV), middle east respiratory syndrome (MERS-CoV), and SARS-CoV-2 viruses are primarily focused ^1–3^. The coronavirus’s structural proteins make up the viral symmetry and enclose the positive-sense single-stranded RNA of ∼30 kb size ^1^. Briefly, the spike protein (S) has S1 and S2 subunits, which recognizes the human receptor ACE-2 and mediates the viral membrane fusion with the host plasma membrane ^4,5^. Whereas, the nucleocapsid protein (N) is phosphorylated and highly basic in nature, whose primarily function is associated with the packaging of viral genomic RNA ^6,7^. The CoV’s N protein contain two RNA-binding domains: the N-terminal domain and the C-terminal domain, linked by a serine/arginine-rich domain (SRD) ^8–11^. The role of SRD is vital for effective virus replication ^12^. In comparison the membrane protein (M) is a transmembrane protein consisting of an N-terminal ectodomain and a C-terminal endodomain ^13–15^.

Viruses use protein-protein interactions (PPI) to reach out and hijack their host cellular network ^16,17^. The virus-host PPI map is invaluable, as it provides insight into the virus’s behavior to capture host protein network for its meanings ^18–20^. Recently, targeting of virus (SARS-CoV-2)-host PPI shows 66 druggable human proteins/host factors targeted by 69 compounds ^16^. Experimental techniques such as biomolecular fluorescence complementation, co-immunoprecipitation, and yeast two-hybrid are extensively used to detect virus-host PPI, which also shed light on the intraviral PPI ^21–24^. The M protein expressed in higher propensity during infection interact with N protein and plays a vital role in assembling virus particles ^25–27^. The M-M interaction occurs by the transmembrane domain ^28^. Further, the N and S proteins interacts with the C-terminal endodomain of M protein, which is the hotspot for protein-protein interaction ^27,29–32^. Besides the role of M protein’s C-terminal in M-N interactions, multiple regions of M protein are responsible for M-E and M-S interactions ^26^. In SARS-CoV, the amino acids 168–208 in the N protein are essential for oligomerization and N-M interactions ^25^. PPI plays a critical role in stabilizing N protein-RNA interactions ^33^. However, the N protein interaction with the C terminal of M protein involves multiple M endodomain regions ^28^. But it is not known in the case of SARS-CoV-2 whether these regions interact or not?

On the other side, computational techniques such as protein-protein interaction networks based on phylogeny methods and structure-based protein-protein docking are now very impactful and faster to identify the interaction sites in protein ^34,35^. In this context, we propose to study the protein-protein interaction of M-E, M-S, and M-N of SARS-CoV-2 with protein-protein docking and molecular dynamics (MD) simulation methods. The primary goal of performing docking is to reveal interaction sites and the generation of protein-protein complexes. Further, atomic-level MD simulations help to characterize the structure and dynamics of protein-protein complexes ^36^. In this study, MD allows us to understand the association-dissociation propensity of protein complex during a single trajectory. Moreover, the study’s outcome will highlight the mechanistic details, i.e., intermediates and transition state, along with the protein complex’s association-dissociation, which could be used as a potential drug target to counter the pathogenicity associated with SARS-CoV-2.

## Material and Methods

### Protein structure modeling and preparation

Many SARS-CoV-2 proteins structure, i.e., spike, protease, and RdRp, are reported by X-ray crystallography or Cryo-EM techniques ^37–39^. However, several other proteins, such as full-length nucleocapsid, envelope, and membrane, do not have structure available yet. Therefore, we have utilized the structure models of the envelope, and membrane proteins using RaptorX web server. We also built the model for the full-length 3D structure of S protein using the I-Tasser web server ^40^, by applying existing Spike protein structures such as PDB ID: 6VXX as template, as the available 3D structures of S protein lack transmembrane and cytosolic part and used it for protein-protein docking. Moreover, the available 3D structures of spike (PDB ID: 6VXX) and envelope (PDB ID: 7K3G) proteins were retrieved from RCSB-PDB for truncated structure docking. The S protein structure is determined using electron microscopy in closed state formed by three S protein monomers. Whereas, the E protein structure is determined in its pentameric form which only constitutes its transmembrane regions in pentameric form. In last, all protein structures were prepared using the protein preparation wizard for optimizing hydrogens and minimizing potential energy using our previously defined protocols ^41,42^.

### Protein-protein docking

The PIPER program embedded in the BioLuminate module of Schrodinger for protein-protein docking was implemented to docking of M protein with E, S, and N proteins ^43,44^. A detailed methodology has been given in our previous report ^41^. Briefly, PIPER performs a global search with Fast-Fourier Transform (FFT) approach and reduces the false-positive results. Among 1000 conformations of input structures, the top 50 clusters were selected with a cluster radius of 9 Å. The docking outcomes based on cluster size were evaluated. With the most massive cluster size, the docked complex out of 5 complexes was selected for molecular dynamics simulation. A total of 70,000 rotations were allowed to generate five docked complexes for all setups.

### MD Simulations of protein-protein complexes

For MD simulation of docked protein-protein complex, three setups were generated for M-E, M-N, and M-S proteins. The binding and their interacting stability were observed for a 100 ns timescale. Using our previously reported protocols, simulation of these complexes carried in the Desmond simulation package, which utilizes OPLS 2005 forcefield to calculate bonded and non-bonded parameters and energy parameters ^45,46^. Previously, the C-terminal region of SARS-CoV M protein was found to interact with N protein in presence of lipids ^26^. Therefore, in our study, simulation of the M-N protein complex was provided with a lipid bilayer (POPE; 1-Palmitoyl-2-oleoyl-sn-glycero-3-phosphoethanolamine) environment around M’s transmembrane regions (residues 20-40, 51-71, and 80-100). All systems fed up with the TIP4P water model, 0.15 M NaCl salt, neutralizing counterions, and minimized for 5000 iterations using the steepest descent method. Final production run, followed by equilibration with NPT ensemble, carried out at an average temperature of 310K, and 1 bar pressure maintained using Noose-Hover chain thermostat and Martyna-Tobias-Klein barostat methods.

### Prime MM-GBSA: Binding energy calculation

Prime module of Schrodinger suite was utilized to calculate binding energy of every protein-protein complex by keeping membrane as a receptor and other three proteins envelope, spike, and nucleocapsid as ligands using VSGB solvation model and OPLS 2005 forcefield. Prime energy and MM-GBSA scores are calculated which refers to the contribution of covalent interaction in the complex and binding energy of protein-protein complex, respectively.

## Results

### Membrane-Envelope interaction

As shown in **figure 1**, the protein-protein complex of M and E proteins have been formed by multiple aromatic hydrogen bonds and a π-cation bond through residues Leu51, Thr55, Phe96, and Phe103 towards the N-terminal of Membrane protein (**Figure 1 and Table 1**). The binding energy calculated for M-E docked complex from the Prime module was found to be 38.96 kcal/mol. On the other hand, the prime energy calculation showed high contribution of covalent interactions with a score of −10906 kcal/mol. Further, the complex was subjected to MD simulations for 100 ns and analyzed for its stability (**Supplementary movie 1**). We have also calculated the simulated frames’ binding energy at every 25 ns of the trajectory (**Figure 5 and Supplementary Table 1**). Additionally, the frames at every 25 ns interval are shown in **figure 2A-2D** and detailed interaction analysis of all four captured snapshots are tabulated in **Supplementary Table 2**, where multiple residues of M protein such as Asp3, Phe45, Trp55, Phe96, Tyr178, etc. are in contact throughout with the E protein through multiple interactions which makes it a stable complex.

**Figure 1:**
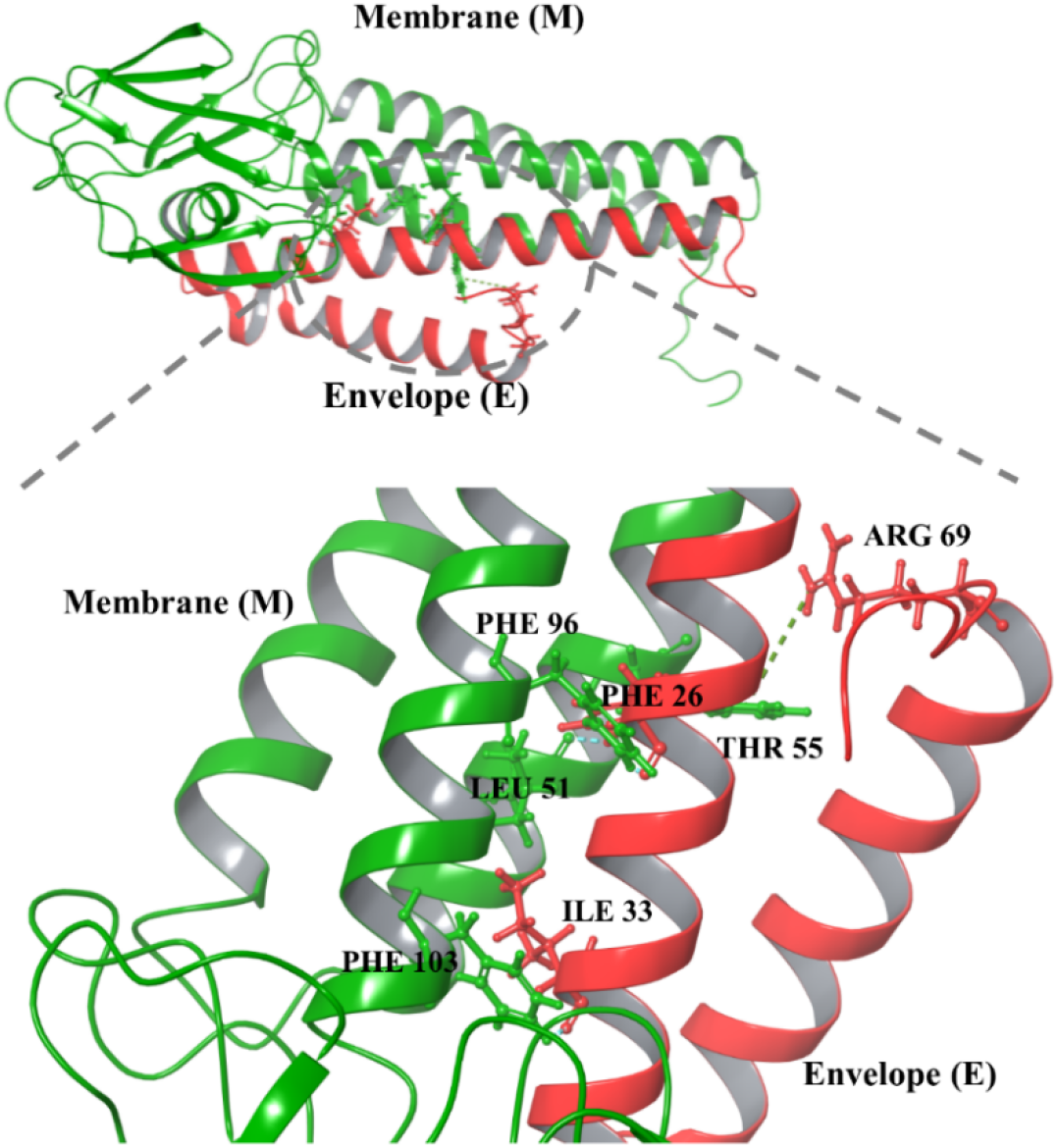
Protein-protein docking of M and E proteins structure models. The colored dashed lines represent the interactions and interacting residues highlighted with ball and stick form in different colors (green of membrane and red of envelope proteins).

**Table 1:** Interaction analysis of protein-protein complexes from computational docking.

| Proteins | Membrane Residue | Envelope Residue | Interaction Type |
| --- | --- | --- | --- |
| **Membrane-Envelope** | LEU 51 | PHE 26 | Aromatic H-bond |
| | THR 55 | ARG 69 | π-cation |
|  | PHE 96 | PHE 26 | Aromatic H-bond |
|  | PHE 103 | ILE 33 | Aromatic H-bond |
| **Membrane-Spike** | **Membrane Residue** | **Spike Residue** |  |
|  | MET 1 | MET 1 | H-bond |
| | MET 1 | PHE 4 | π-cation |
|  | ASN 5 | GLN 173 | H-bond |
|  | TYR 71 | ASP 198 | H-bond |
| | TRP 75 | PHE 377 | π-π stacking |
|  | ARG 174 | HIS 625 | Aromatic H-bond |
| **Membrane-Nucleocapsid** | **Membrane Residue** | **Nucleocapsid Residue** |  |
|  | TYR 199 | GLY 335 | H-bond |
|  | ASP 209 | PHE 314 | Aromatic H-bond |
|  | HIS 210 | PHE 286 | Aromatic H-bond |

**Figure 2:**
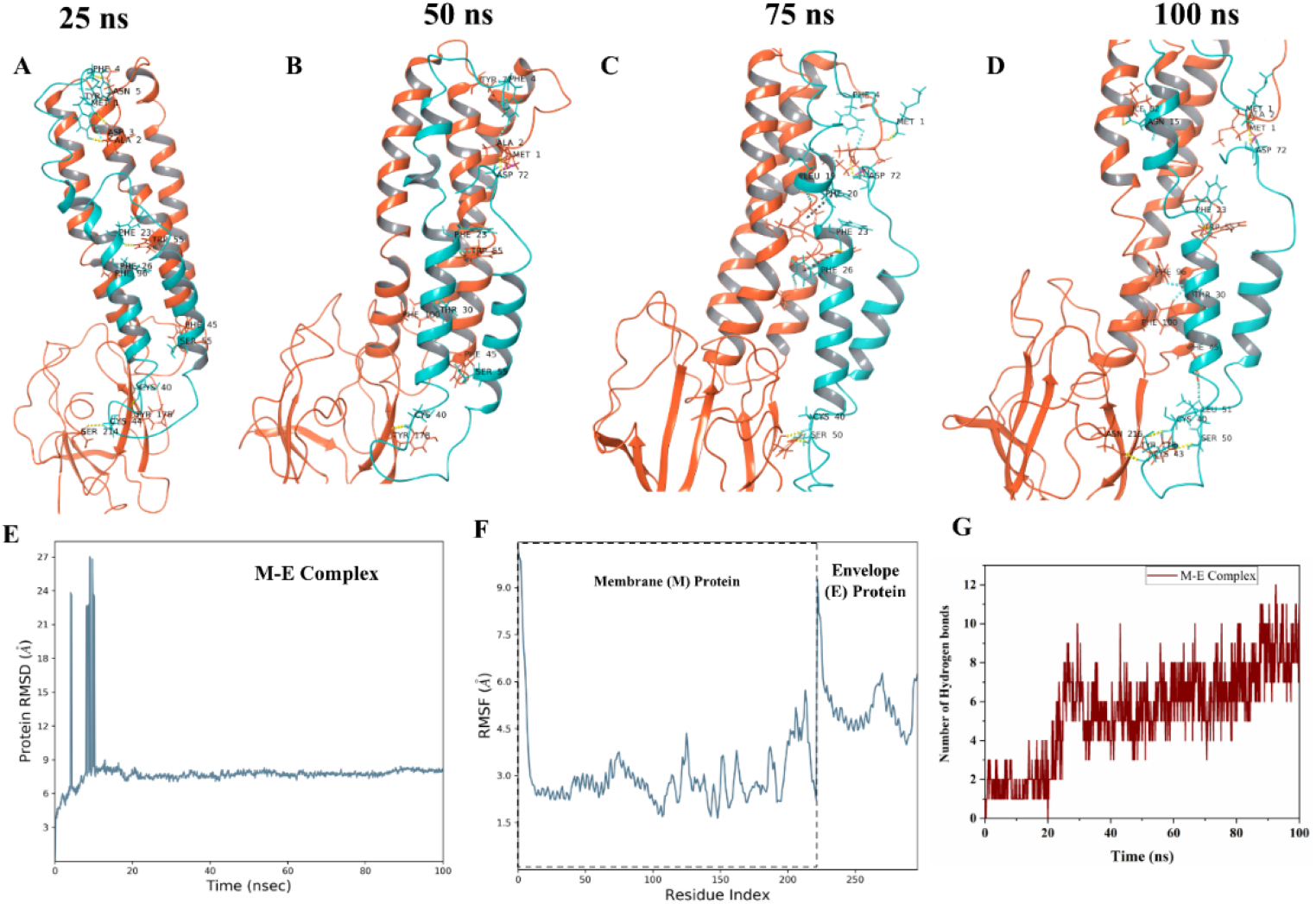
Molecular dynamics simulations of M-E docked complex. **A-D.** The snapshots at every 25 ns interval of 100 ns simulation trajectory of interacting complex, highlighting all bonded residues. The colored dashed lines represent the interactions and interacting residues highlighted with ball and stick form in different colors (orange of membrane and cyan of envelope proteins). **E.** Root Mean Square Deviation (RMSD) of the complex. **F**. Root Mean Square Fluctuations (RMSF) of both proteins. The dashed line box shows the boundary of membrane and envelope protein residues in the plot, and **G.** Depiction of hydrogen bonds formed between these two proteins.

We have observed high binding energy for the M-E complex after simulation i.e., energy from positive to negative scores shows the change in interaction reaction from non-spontaneous to spontaneous. A gradual decrease of ∼20 kcal/mol in prime energy was also observed at regular time interval frame. As shown in **figure 2**, the M-E complex had shown heavy fluctuations in initial frames till 20 ns but found to be relatively stable with RMSD at ∼8Å throughout rest of the simulation period **(Figure 2E)**. The mean changes of M and E protein residues fluctuations within the interaction site were significantly more compared to the non-interacting region **(Figure 2F)**. Further, the number of hydrogen bonds found increased between both proteins throughout the simulation period, with an average of ∼5 **(Figure 2G)**.

### Membrane-Spike interaction

The S protein interact with M in ERGIC, therefore, these two proteins’ docked complex showed promising interactions viz. multiple hydrogen bonds and an aromatic hydrogen bond, π-cation, and π-π stacking, each. The interacting residues of S proteins were found at C-terminal (**figure 3, table 1**).

**Figure 3:**
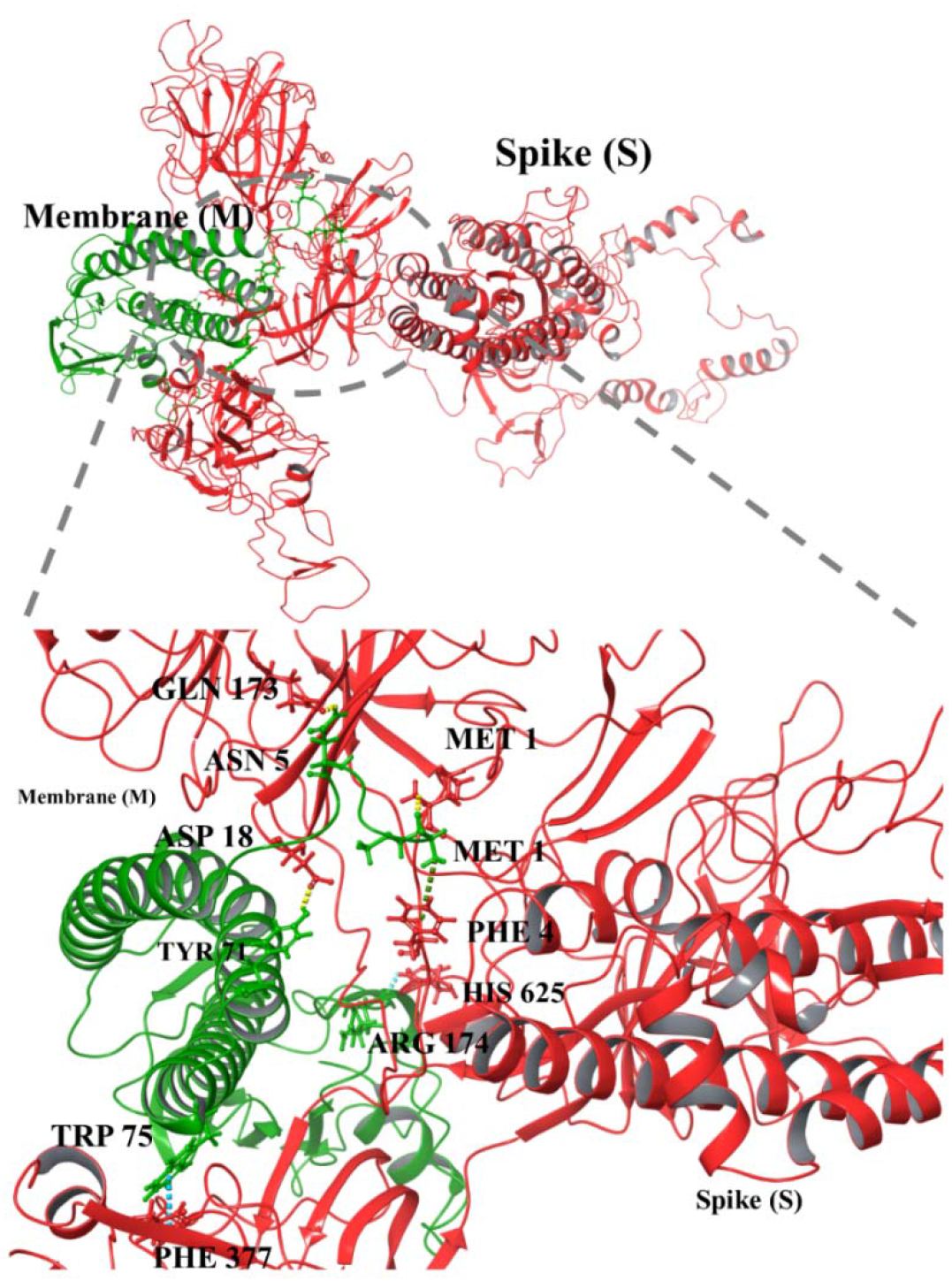
Protein-protein docking of M and S proteins structure models. The ball and stick represent th interacting residues in different colors (green of membrane and red of spike).

The binding energy of the M-S docked complex was calculated to be high, whereas the prime energy calculated to be −49369.2 kcal/mol. We have further investigated the M-S complex’s binding stability through MD simulations up to 100 ns **(Supplementary movie 2)**. From the trajectory, the snapshots at every 25 ns are shown with interacting residues of both proteins which demonstrate that mostly interacting residues are retained during simulations **(Figure 4A-4D)** and their interactions are illustrated in **Supplementary Table 3**. As per the interaction analysis, the residues of M protein such as Asn5, Phe96, Arg174, Asn207 are interacting with S protein at multiple regions constantly with multiple strong non-covalent interactions throughout the simulations. The RMSD values from MD simulation trajectory were trending upward from 5 to 18 Å with a little stabilized trajectory in the entire simulation period (**Figure 4E**). The RMSF plot of the loosely packed S protein model with 1273 residues showed massive fluctuations near 630^th^-750^th^ residues up to 18Å (**Figure 4F**). However, the fluctuations in interacting site residues of S protein’s C-terminal were around 18Å. The binding free energy from the simulation trajectory of M-S complexes is represented in **Figure 5 (**and tabulated in **Supplementary Table 1)** which has shown a constant decrease in positive binding energy and a gradual increase in prime energy. In final, the average number of hydrogen bonds were ∼12 in M-S complex simulation setup throughout the MD period (**Figure 4G**).

**Figure 4:**
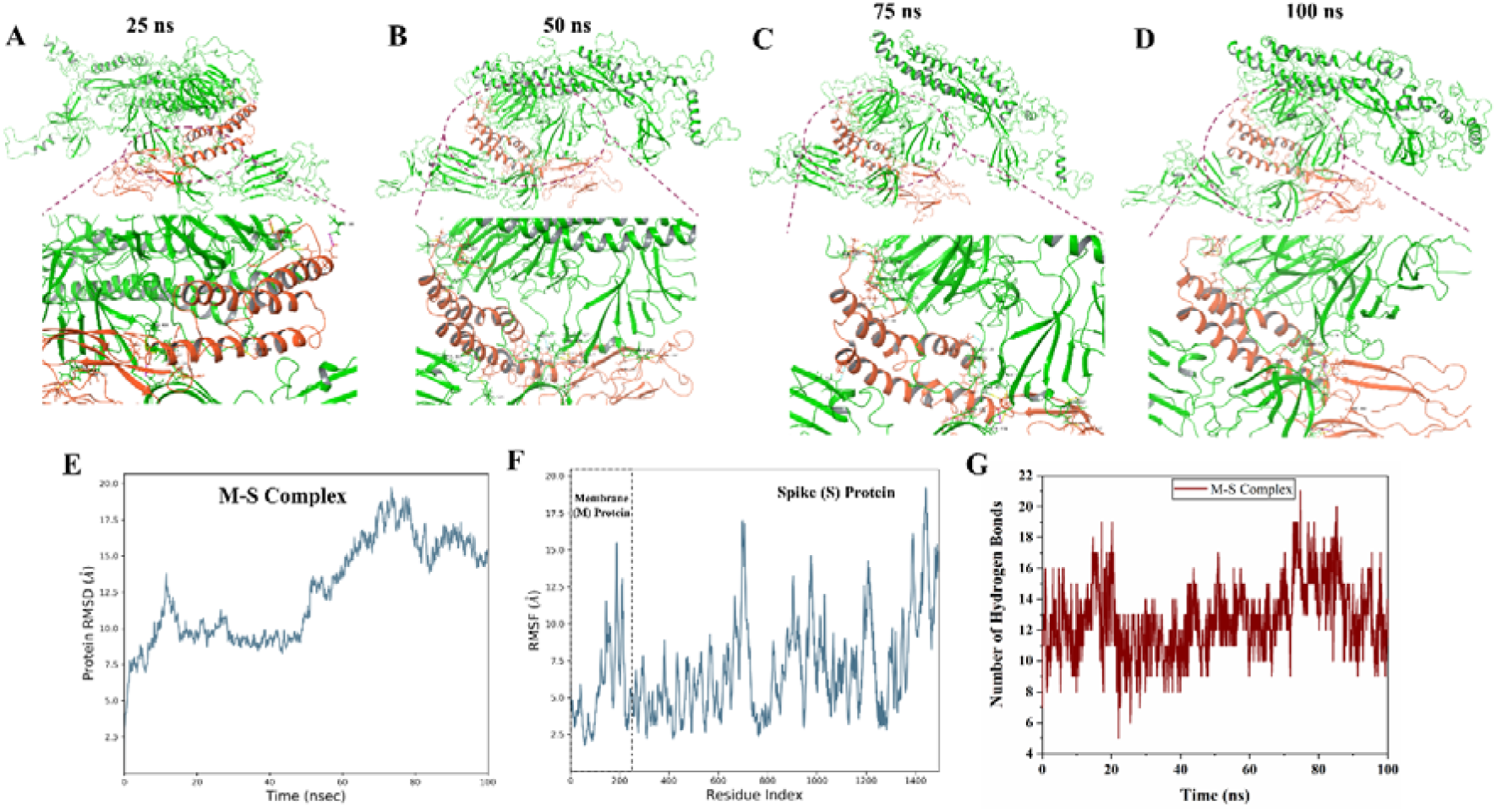
Molecular dynamics simulations of M-S docked complex. **A-D.** The snapshots at every 25 ns interval of 100 ns simulation trajectory of interacting complex, highlighting all bonded residues. The colored dashed lines represent the interactions and interacting residues highlighted with ball and stick form in different colors (orange of membrane and green of spike proteins). **E.** Root Mean Square Deviation (RMSD) of the complex. **F**. Root Mean Square Fluctuations (RMSF) of both proteins. Th dashed line box shows the boundary of membrane and spike protein residues in the plot, and **G.** Depiction of hydrogen bonds formed between these two proteins.

**Figure 5:**
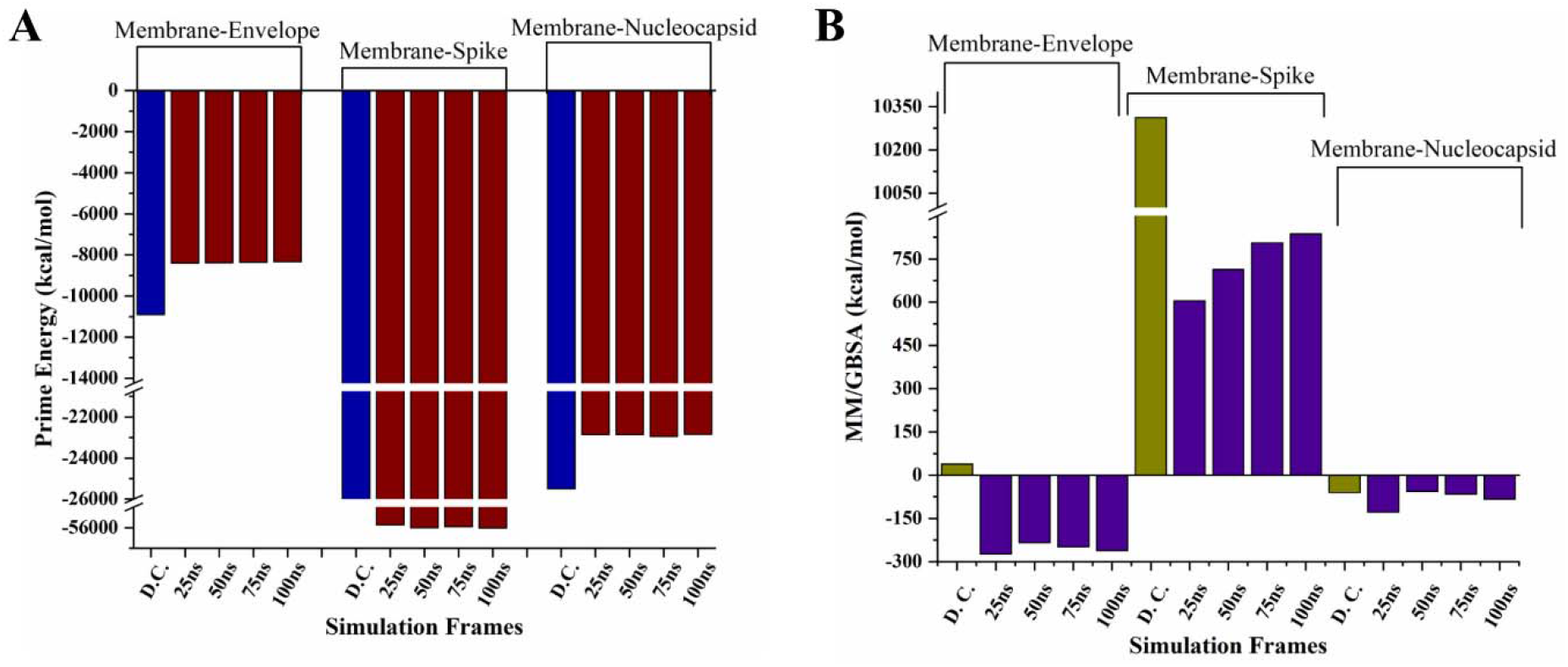
Prime energy (A) and Binding energy (B) calculation of protein-protein complexes using MM-GBSA approach. The complexes are selected at every 25ns of simulation trajectory and compared with the docked complex (obtained from protein-protein docking).

### Membrane-Nucleocapsid interaction

The protein-protein docking of M-N complex showed a total of three residues of M protein viz. Tyr199, Asp209, and His210 are interacting with residues Gly335, Phe314, and Phe286 of N protein via one hydrogen bond and two aromatic hydrogen bonds, respectively (**Figure 6 and Table 1**). Moreover, the docked complex M-N has attained the high binding energy of −59.8 kcal/mol (**Figure 5 and Supplementary Table 1**).

**Figure 6:**
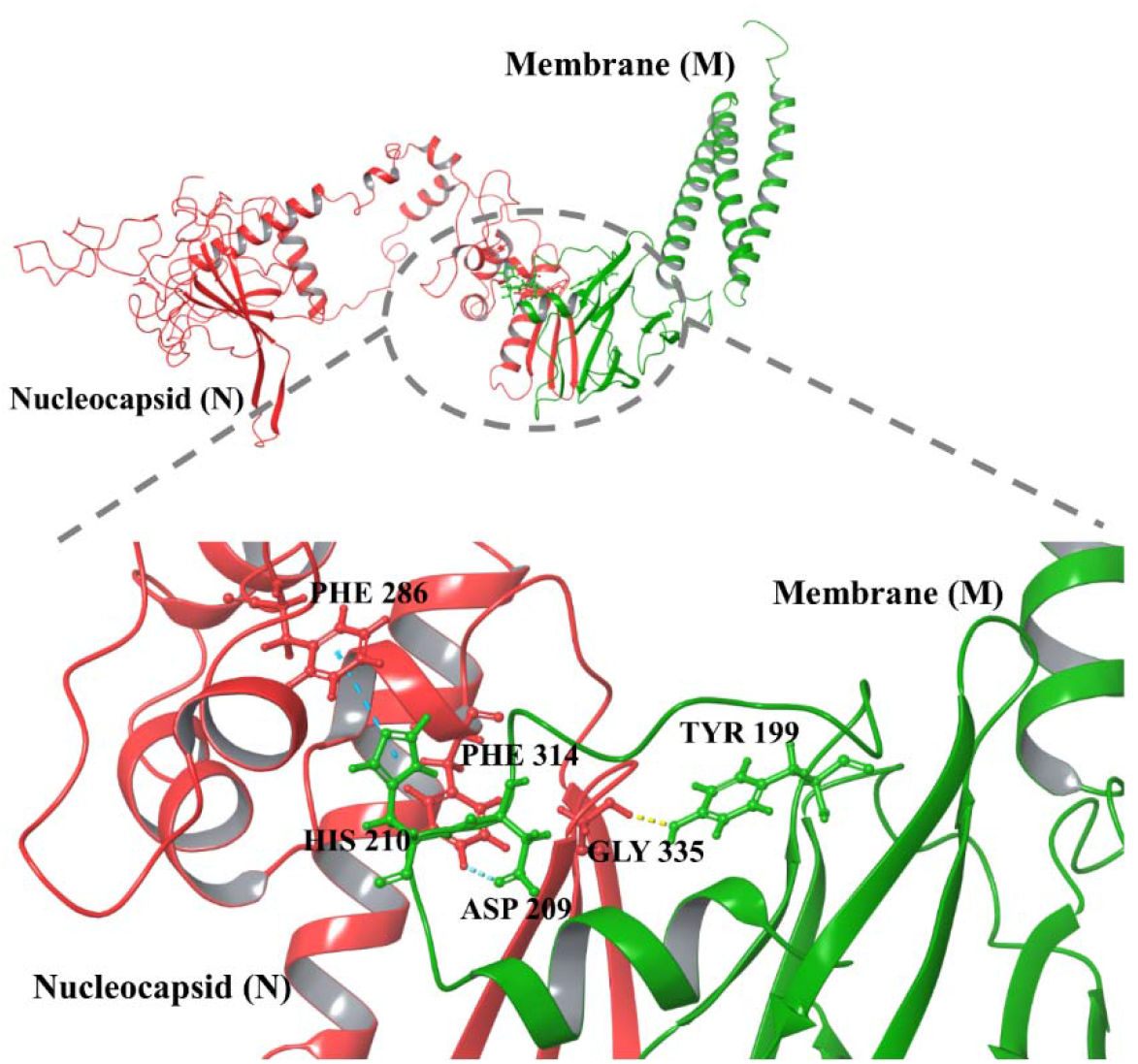
Protein-protein docking of M and N protein structure models. The ball and stick represent interacting residues in different colors (green of Membrane and red of Nucleocapsid).

Further, the slight changes in interacting residues of the complex are shown in snapshots from the 100 ns simulation trajectory at every 25 ns (**Figure 7A-7D**; see **Supplementary Table 4** for residue interactions). The residues such as Arg150, Asn207, etc. of M protein contribute to the contact establishment with N protein’s residues during simulation period. The M-N protein-protein complex was observed with an average RMSD of approx. 11 Å based on simulation analysis **(Figure 7E)**. However, there was a fluctuating trend in RMSF values throughout the simulation from 2Å to 6Å in N protein residues. These fluctuations may be due to high disorder propensity in N protein and can be seen in the **Supplementary movie 3**. The RMSF values of interacting residues of M protein were 1.7 Å (Trp58), 1.2 Å (Arg107), 2.1 Å (Asp163) and for N protein 4.9 Å (Lys256), 2.2 Å (Ser184), and 2.9 Å (Tyr268) for 100 ns simulation period **(Figure 7F)**. The number of intermediate hydrogen bonds formed within the simulation setup was ∼ 7 up to 100 ns timescale **(Figure 7G)**. The binding free energy of complexes from the simulation trajectory is higher than the complex (except the frame at 50 ns) obtained from protein-protein docking **(Figure 5 and Supplementary Table 1)**.

**Figure 7:**
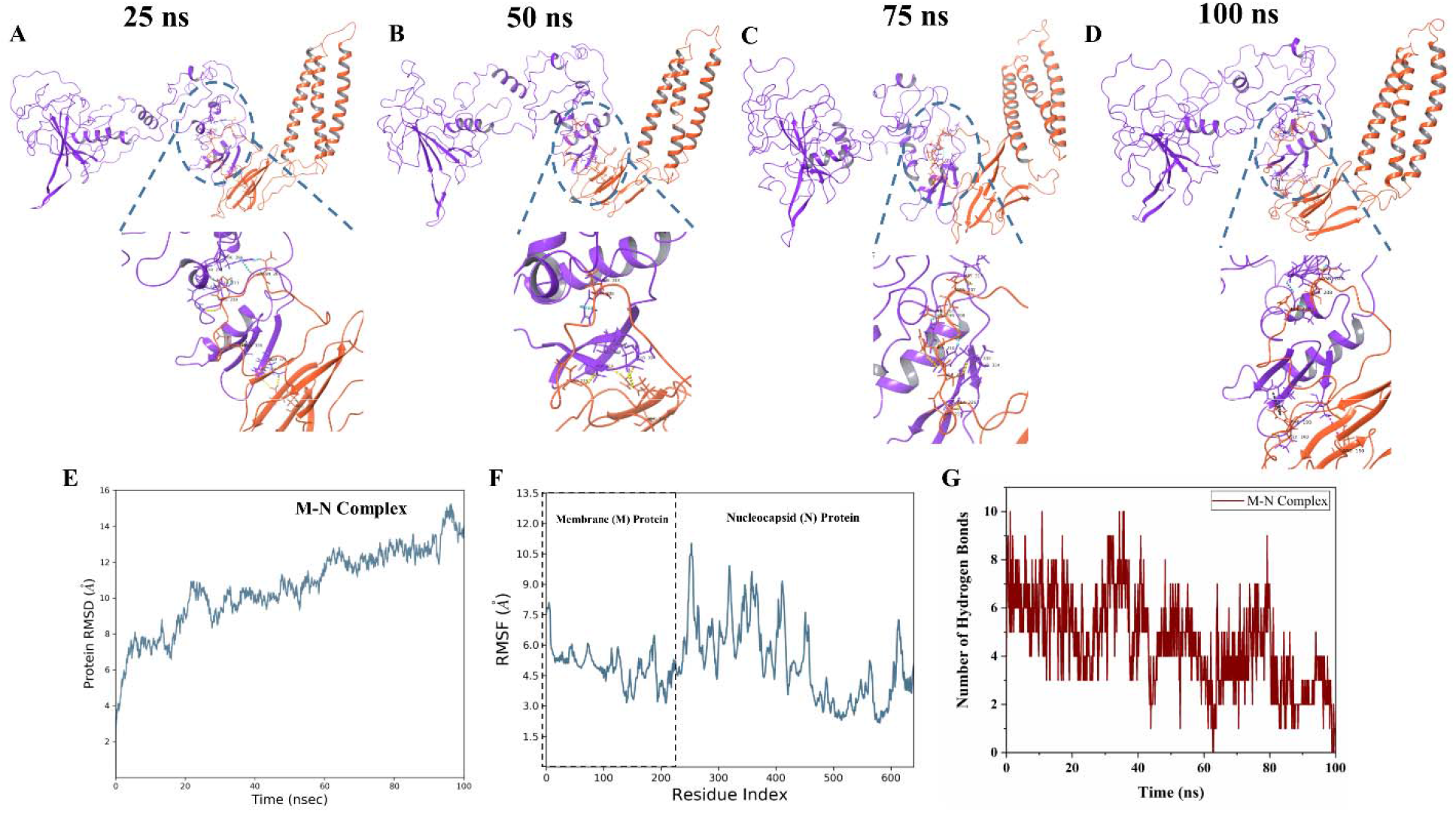
Molecular dynamics simulations of M-N docked complex. **A-D.** The snapshots at every 25 ns interval of 100 ns simulation trajectory of interacting complex, highlighting all bonded residues. The colored dashed lines represent the interactions and interacting residues highlighted with ball and stick form in different colors (orange of membrane and violet of nucleocapsid proteins). **E.** Root Mean Square Deviation (RMSD) of the complex. **F**. Root Mean Square Fluctuations (RMSF) of both proteins. The dashed line box shows the boundary of membrane and nucleocapsid protein residues in the plot, and **G.** Depiction of hydrogen bonds formed between these two proteins.

## Discussion

Intraviral Protein-Protein interactions play an essential role in the coronavirus life cycle, specifically during the replicating complex formation, as elucidated from several structural studies ^47–49^. The RNA dependent RNA polymerase (nsp12) of SARS-CoV interacts with nsp7 and nsp8 and increases the RNA-synthesizing activity ^47^. The nsp12-nsp7-nsp8 also associate with the nsp14 (proofreading enzyme) ^47^. The cryo-EM studies showed that the nsp7 and nsp8 heterodimers stabilize RNA binding regions of nsp12, while the second subunit of nsp8 plays a vital role in polymerase activity ^48^. Further, structural studies showed that nsp10 interacts with the N-terminal domain of nsp14 to stabilize it and stimulate its activity ^49^.

Similarly, the SARS-CoV structural proteins have been reported to interact with each other and play an essential role in virus assembly ^6,15,28^. Therefore, in this study, we report the intraviral PPI among structural proteins of SARS-CoV-2. We have computationally shown that the M proteins interact with other structural proteins to form complexes of M-E, M-S, and M-N, responsible for the proper virus assembly. We have performed protein-protein docking to identify the regions and residues which interact during these bindings. We have investigated these in membrane protein with several interacting structural proteins such as envelope, spike, and nucleocapsid proteins, respectively. Previously, in SARS-CoV, mutation-based studies showed that M protein is vital for virus assembly and interact with other structural proteins ^26^. The entire C-terminus domain of M proteins was found to interact with N protein ^26,29,31^. Similarly, two transmembrane domains and the cytoplasmic domain of M protein were reported to interact with E protein ^26^. There are multiple regions of M protein that interact with spike glycoprotein ^26^.

Therefore, in this study, we have considered the M protein as a receptor and S, E, and N proteins as protein ligands. We also checked the interaction of M protein with a truncated structure (residues 8-38) of E protein which is in its pentameric form and the interacting residues are shown in **Supplementary Figure 1**. As revealed in this study, multiple regions of S interacts with M protein, which has also been seen in other coronaviruses ^50^. The S protein exists in its trimeric form therefore, we have also docked the trimeric crystal structure (PDB ID: 6VXX) with M protein, where, few interactions which include one hydrogen bond and four π-π stacking were observed **(Supplementary Figure 2)**. The M protein is a triple-spanning membrane protein. Its cytosolic region is solely responsible for M-N interaction; therefore, in the case of M-N docking, the cytosolic part of M was targeted for interaction with N protein.

To understand the stability of docked complexes and formed interactions, we have performed 100 ns long MD simulations. The simulation studies showed resilience in docked protein complexes of M-E, M-S, and M-N. The binding energy was found in good agreement with the results and allowed good binding of intraviral structural proteins. Our computational studies agree with previous reports, where particle assembly occurs in the Endoplasmic Reticulum-Golgi intermediate compartment (ERGIC) and finally trafficked for release via exocytosis ^51^ (**Figure 8)**

**Figure 8.**
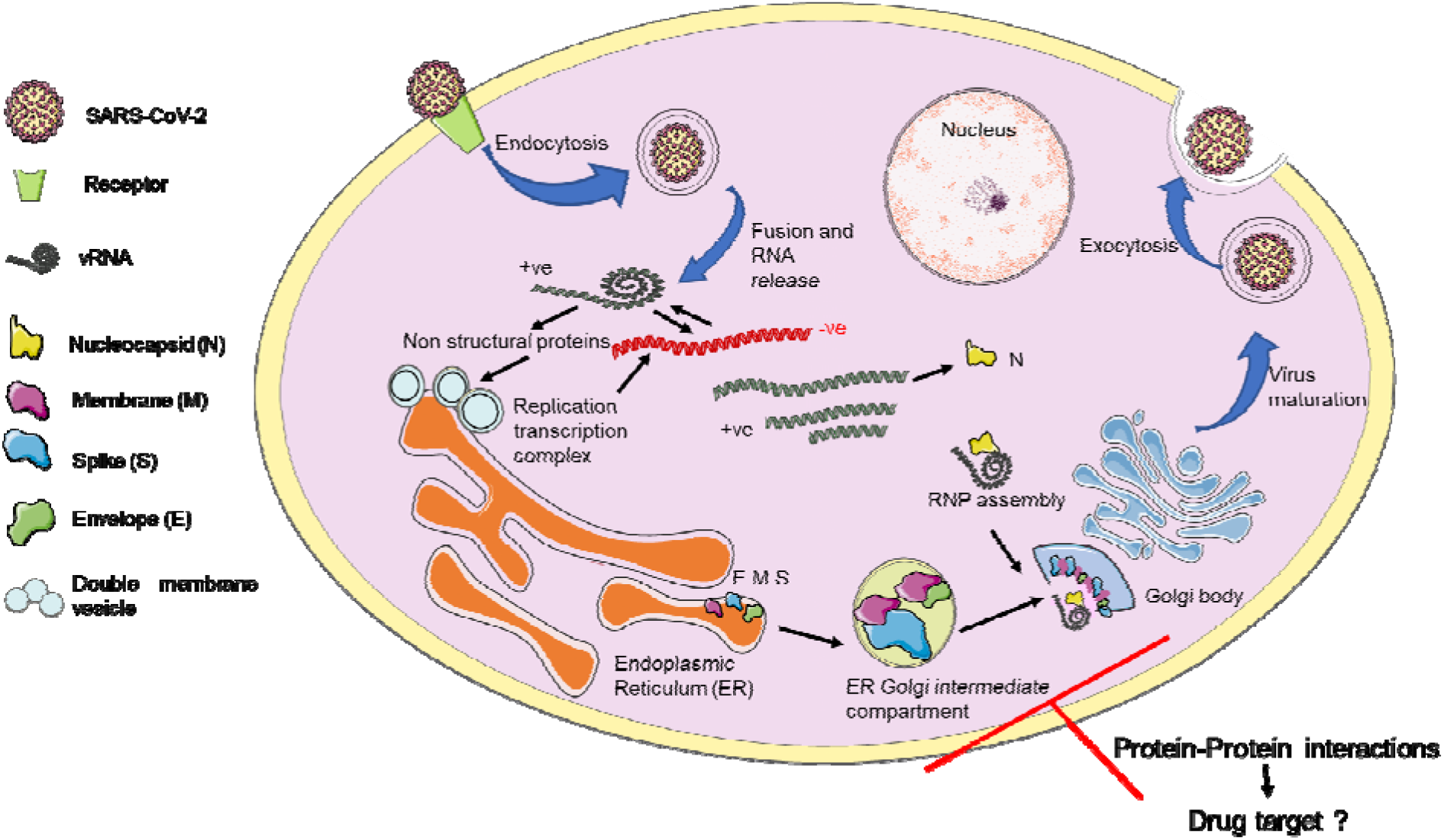
Schematic representation of protein-protein interactions among SARS-CoV-2 structural proteins (Membrane, Spike, Nucleocapsid, and Envelope).

## Conclusion

Despite the small genome of viruses, they are highly pathogenic/infectious, and their genome integrity allows them to hijack the host cellular machinery. For rapid infection and replication, viruses follow multiple pathways. In between regulating the host cellular system, it is essential to coordinate among own proteins for proper assembly and genome encapsulation. Here, PPI plays an essential role in coronaviruses where structural protein interacts with each other, encapsulate the genome, and forms mature viruses. It could be a great interest to study these PPIs in drug targeting, as disruption of virus assembly will lead to immature virion formation. In this context, the present study may help to design the mutation-based studies to understand PPI in SARS-CoV-2 and targeting several interacting residues for therapeutic purposes. Also, it would be interesting to investigate these structural proteins’ interaction specifically with several host proteins. Moreover, the driving forces which lead to the formation of proteins assembly and virus particle formations could also be examined. Additional studies on binding mechanism and energy favorable interaction of structural protein could help us in developing new strategies against protein-protein interaction.

## Author Contribution

RG, NG: study supervision and designed the experiment. AK and PK acquisition and interpretation of computational data. AK, PK, and RG contributed to paper writing. PK and AK have contributed equally.

## Declaration of competing interest

All authors affirm that there are no conflicts of interest.

## Acknowledgments

All the authors would like to thank IIT Mandi for the infrastructure. RG is thankful to IYBA award from DBT, Government of India (BT/11/IYBA/2018/06) and SERB, Govt. Of India (CRG/2019/005603).

## Conflict of Interest

All authors affirm that there are no conflicts of interest.

## Notes

### Competing Interest Statement

The authors have declared no competing interest.

## References

(1) Giri, R.; Bhardwaj, T.; Shegane, M.; Gehi, B. R.; Kumar, P.; Gadhave, K.; Oldfield, C. J.; Uversky, V. N. Understanding COVID-19 via Comparative Analysis of Dark Proteomes of SARS-CoV-2, Human SARS and Bat SARS-like Coronaviruses. Cell. Mol. Life Sci. 2020. https://doi.org/10.1007/s00018-020-03603-x.

(2) Kumar, A.; Kumar, A.; Kumar, P.; Garg, N.; Giri, R. SARS-CoV-2 NSP1 C-Terminal Region (Residues 130-180) Is an Intrinsically Disordered Region. bioRxiv 2020.

(3) Gadhave, K.; Kumar, P.; Kumar, A.; Bhardwaj, T.; Garg, N.; Giri, R. NSP 11 of SARS-CoV-2 Is an Intrinsically Disordered Protein. bioRxiv 2020, 2020.10.07.330068. https://doi.org/10.1101/2020.10.07.330068.

(4) Li, F. Structure, Function, and Evolution of Coronavirus Spike Proteins. Annu. Rev. Virol. 2016, 3(1), 237–261. https://doi.org/10.1146/annurev-virology-110615-042301.

(5) Yan, R.; Zhang, Y.; Li, Y.; Xia, L.; Guo, Y.; Zhou, Q. Structural Basis for the Recognition of SARS-CoV- 2 by Full-Length Human ACE2. Science 2020, 367(6485), 1444–1448. https://doi.org/10.1126/science.abb2762.

(6) He, R.; Dobie, F.; Ballantine, M.; Leeson, A.; Li, Y.; Bastien, N.; Cutts, T.; Andonov, A.; Cao, J.; Booth, T. F.; Plummer, F. A.; Tyler, S.; Baker, L.; Li, X. Analysis of Multimerization of the SARS Coronavirus Nucleocapsid Protein. Biochem. Biophys. Res. Commun. 2004, 316(2), 476–483. https://doi.org/10.1016/j.bbrc.2004.02.074.

(7) Zeng, W.; Liu, G.; Ma, H.; Zhao, D.; Yang, Y.; Liu, M.; Mohammed, A.; Zhao, C.; Yang, Y.; Xie, J.; Ding, C.; Ma, X.; Weng, J.; Gao, Y.; He, H.; Jin, T. Biochemical Characterization of SARS-CoV-2 Nucleocapsid Protein. Biochem. Biophys. Res. Commun. 2020, 527(3), 618–623. https://doi.org/10.1016/j.bbrc.2020.04.136.

(8) Huang, Q.; Yu, L.; Petros, A. M.; Gunasekera, A.; Liu, Z.; Xu, N.; Hajduk, P.; Mack, J.; Fesik, S. W.; Olejniczak, E. T. Structure of the N-Terminal RNA-Binding Domain of the SARS CoV Nucleocapsid Protein. Biochemistry 2004, 43(20), 6059–6063. https://doi.org/10.1021/bi036155b.

(9) Luo, H.; Chen, J.; Chen, K.; Shen, X.; Jiang, H. Carboxyl Terminus of Severe Acute Respiratory Syndrome Coronavirus Nucleocapsid Protein:⍰ Self-Association Analysis and Nucleic Acid Binding Characterization. Biochemistry 2006, 45(39), 11827–11835. https://doi.org/10.1021/bi0609319.

(10) Luo, H.; Ye, F.; Chen, K.; Shen, X.; Jiang, H. SR-Rich Motif Plays a Pivotal Role in Recombinant SARS Coronavirus Nucleocapsid Protein Multimerization. Biochemistry 2005, 44(46), 15351–15358. https://doi.org/10.1021/bi051122c.

(11) Cubuk, J.; Alston, J. J.; Incicco, J. J.; Singh, S.; Stuchell-Brereton, M. D.; Ward, M. D.; Zimmerman, M. I.; Vithani, N.; Griffith, D.; Wagoner, J. A.; Bowman, G. R.; Hall, K. B.; Soranno, A.; Holehouse, A. S. The SARS-CoV-2 Nucleocapsid Protein Is Dynamic, Disordered, and Phase Separates with RNA. bioRxiv 2020. https://doi.org/10.1101/2020.06.17.158121.

(12) Tylor, S.; Andonov, A.; Cutts, T.; Cao, J.; Grudesky, E.; Van Domselaar, G.; Li, X.; He, R. The SR-Rich Motif in SARS-CoV Nucleocapsid Protein Is Important for Virus Replication. Can. J. Microbiol. 2009, 55(3), 254–260. https://doi.org/10.1139/w08-139.

(13) Bianchi, M.; Benvenuto, D.; Giovanetti, M.; Angeletti, S.; Ciccozzi, M.; Pascarella, S. Sars-CoV-2 Envelope and Membrane Proteins: Structural Differences Linked to Virus Characteristics? https://www.hindawi.com/journals/bmri/2020/4389089/ (accessed 2020 -10 -09). https://doi.org/10.1155/2020/4389089.

(14) Rottier, P.; Brandenburg, D.; Armstrong, J.; van der Zeijst, B.; Warren, G. Assembly in Vitro of a Spanning Membrane Protein of the Endoplasmic Reticulum: The E1 Glycoprotein of Coronavirus Mouse Hepatitis Virus A59. Proc. Natl. Acad. Sci. U. S. A. 1984, 81(5), 1421–1425. https://doi.org/10.1073/pnas.81.5.1421.

(15) Artika, I. M.; Dewantari, A. K.; Wiyatno, A. Molecular Biology of Coronaviruses: Current Knowledge. Heliyon 2020, 6(8), e04743. https://doi.org/10.1016/j.heliyon.2020.e04743.

(16) Gordon, D. E.; Jang, G. M.; Bouhaddou, M.; Xu, J.; Obernier, K.; White, K. M.; O’Meara, M. J.; Rezelj, V. V.; Guo, J. Z.; Swaney, D. L.; Tummino, T. A.; Hüttenhain, R.; Kaake, R. M.; Richards, A. L.; Tutuncuoglu, B.; Foussard, H.; Batra, J.; Haas, K.; Modak, M.; Kim, M.; Haas, P.; Polacco, B. J.; Braberg, H.; Fabius, J. M.; Eckhardt, M.; Soucheray, M.; Bennett, M. J.; Cakir, M.; McGregor, M. J.; Li, Q.; Meyer, B.; Roesch, F.; Vallet, T.; Mac Kain, A.; Miorin, L.; Moreno, E.; Naing, Z. Z. C.; Zhou, Y.; Peng, S.; Shi, Y.; Zhang, Z.; Shen, W.; Kirby, I. T.; Melnyk, J. E.; Chorba, J. S.; Lou, K.; Dai, S. A.; Barrio-Hernandez, I.; Memon, D.; Hernandez-Armenta, C.; Lyu, J.; Mathy, C. J. P.; Perica, T.; Pilla, K. B.; Ganesan, S. J.; Saltzberg, D. J.; Rakesh, R.; Liu, X.; Rosenthal, S. B.; Calviello, L.; Venkataramanan, S.; Liboy-Lugo, J.; Lin, Y.; Huang, X.-P.; Liu, Y.; Wankowicz, S. A.; Bohn, M.; Safari, M.; Ugur, F. S.; Koh, C.; Savar, N. S.; Tran, Q. D.; Shengjuler, D.; Fletcher, S. J.; O’Neal, M. C.; Cai, Y.; Chang, J. C. J.; Broadhurst, D. J.; Klippsten, S.; Sharp, P. P.; Wenzell, N. A.; Kuzuoglu-Ozturk, D.; Wang, H.-Y.; Trenker, R.; Young, J. M.; Cavero, D. A.; Hiatt, J.; Roth, T. L.; Rathore, U.; Subramanian, A.; Noack, J.; Hubert, M.; Stroud, R. M.; Frankel, A. D.; Rosenberg, O. S.; Verba, K. A.; Agard, D. A.; Ott, M.; Emerman, M.; Jura, N.; von Zastrow, M.; Verdin, E.; Ashworth, A.; Schwartz, O.; d’Enfert, C.; Mukherjee, S.; Jacobson, M.; Malik, H. S.; Fujimori, D. G.; Ideker, T.; Craik, C. S.; Floor, S. N.; Fraser, J. S.; Gross, J. D.; Sali, A.; Roth, B. L.; Ruggero, D.; Taunton, J.; Kortemme, T.; Beltrao, P.; Vignuzzi, M.; García-Sastre, A.; Shokat, K. M.; Shoichet, B. K.; Krogan, N. J. A SARS-CoV-2 Protein Interaction Map Reveals Targets for Drug Repurposing. Nature 2020, 583(7816), 459–468. https://doi.org/10.1038/s41586-020-2286-9.

(17) Kumar, A.; Kumar, P.; Giri, R. Zika Virus NS4A Cytosolic Region (Residues 1–48) Is an Intrinsically Disordered Domain and Folds upon Binding to Lipids. Virology 2020, 550, 27–36. https://doi.org/10.1016/j.virol.2020.07.017.

(18) Brito, A. F.; Pinney, J. W. Protein–Protein Interactions in Virus–Host Systems. Front. Microbiol. 2017, 8. https://doi.org/10.3389/fmicb.2017.01557.

(19) Lasso, G.; Mayer, S. V.; Winkelmann, E. R.; Chu, T.; Elliot, O.; Patino-Galindo, J. A.; Park, K.; Rabadan, R.; Honig, B.; Shapira, S. D. A Structure-Informed Atlas of Human-Virus Interactions. Cell 2019, 178(6), 1526–1541.e16. https://doi.org/10.1016/j.cell.2019.08.005.

(20) Gordon, D. E.; Hiatt, J.; Bouhaddou, M.; Rezelj, V. V.; Ulferts, S.; Braberg, H.; Jureka, A. S.; Obernier, K.; Guo, J. Z.; Batra, J.; Kaake, R. M.; Weckstein, A. R.; Owens, T. W.; Gupta, M.; Pourmal, S.; Titus, E. W.; Cakir, M.; Soucheray, M.; McGregor, M.; Cakir, Z.; Jang, G.; O’Meara, M. J.; Tummino, T. A.; Zhang, Z.; Foussard, H.; Rojc, A.; Zhou, Y.; Kuchenov, D.; Hüttenhain, R.; Xu, J.; Eckhardt, M.; Swaney, D. L.; Fabius, J. M.; Ummadi, M.; Tutuncuoglu, B.; Rathore, U.; Modak, M.; Haas, P.; Haas, K. M.; Naing, Z. Z. C.; Pulido, E. H.; Shi, Y.; Barrio-Hernandez, I.; Memon, D.; Petsalaki, E.; Dunham, A.; Marrero, M. C.; Burke, D.; Koh, C.; Vallet, T.; Silvas, J. A.; Azumaya, C. M.; Billesbølle, C.; Brilot, A. F.; Campbell, M. G.; Diallo, A.; Dickinson, M. S.; Diwanji, D.; Herrera, N.; Hoppe, N.; Kratochvil, H. T.; Liu, Y.; Merz, G. E.; Moritz, M.; Nguyen, H. C.; Nowotny, C.; Puchades, C.; Rizo, A. N.; Schulze-Gahmen, U.; Smith, A. M.; Sun, M.; Young, I. D.; Zhao, J.; Asarnow, D.; Biel, J.; Bowen, A.; Braxton, J. R.; Chen, J.; Chio, C. M.; Chio, U. S.; Deshpande, I.; Doan, L.; Faust, B.; Flores, S.; Jin, M.; Kim, K.; Lam, V. L.; Li, F.; Li, J.; Li, Y.-L.; Li, Y.; Liu, X.; Lo, M.; Lopez, K. E.; Melo, A. A.; Moss, F. R.; Nguyen, P.; Paulino, J.; Pawar, K. I.; Peters, J. K.; Pospiech, T. H.; Safari, M.; Sangwan, S.; Schaefer, K.; Thomas, P. V.; Thwin, A. C.; Trenker, R.; Tse, E.; Tsui, T. K. M.; Wang, F.; Whitis, N.; Yu, Z.; Zhang, K.; Zhang, Y.; Zhou, F.; Saltzberg, D.; Consortium12†, Q. S. B.; Hodder, A. J.; Shun-Shion, A. S.; Williams, D. M.; White, K. M.; Rosales, R.; Kehrer, T.; Miorin, L.; Moreno, E.; Patel, A. H.; Rihn, S.; Khalid, M. M.; Vallejo-Gracia, A.; Fozouni, P.; Simoneau, C. R.; Roth, T. L.; Wu, D.; Karim, M. A.; Ghoussaini, M.; Dunham, I.; Berardi, F.; Weigang, S.; Chazal, M.; Park, J.; Logue, J.; McGrath, M.; Weston, S.; Haupt, R.; Hastie, C. J.; Elliott, M.; Brown, F.; Burness, K. A.; Reid, E.; Dorward, M.; Johnson, C.; Wilkinson, S. G.; Geyer, A.; Giesel, D. M.; Baillie, C.; Raggett, S.; Leech, H.; Toth, R.; Goodman, N.; Keough, K. C.; Lind, A. L.; Consortium‡, Z.; Klesh, R. J.; Hemphill, K. R.; Carlson-Stevermer, J.; Oki, J.; Holden, K.; Maures, T.; Pollard, K. S.; Sali, A.; Agard, D. A.; Cheng, Y.; Fraser, J. S.; Frost, A.; Jura, N.; Kortemme, T.; Manglik, A.; Southworth, D. R.; Stroud, R. M.; Alessi, D. R.; Davies, P.; Frieman, M. B.; Ideker, T.; Abate, C.; Jouvenet, N.; Kochs, G.; Shoichet, B.; Ott, M.; Palmarini, M.; Shokat, K. M.; García-Sastre, A.; Rassen, J. A.; Grosse, R.; Rosenberg, O. S.; Verba, K. A.; Basler, C. F.; Vignuzzi, M.; Peden, A. A.; Beltrao, P.; Krogan, N. J. Comparative Host-Coronavirus Protein Interaction Networks Reveal Pan-Viral Disease Mechanisms. Science 2020. https://doi.org/10.1126/science.abe9403.

(21) Khorsand, B.; Savadi, A.; Naghibzadeh, M. SARS-CoV-2-Human Protein-Protein Interaction Network. Inform. Med. Unlocked 2020, 20, 100413. https://doi.org/10.1016/j.imu.2020.100413.

(22) Wiederschain, G. Ya. Protein-Protein Interactions. A Molecular Cloning Manual. Biochem. Mosc. 2006, 71(6), 697–697. https://doi.org/10.1134/S0006297906060162.

(23) von Brunn, A.; Teepe, C.; Simpson, J. C.; Pepperkok, R.; Friedel, C. C.; Zimmer, R.; Roberts, R.; Baric, R.; Haas, J. Analysis of Intraviral Protein-Protein Interactions of the SARS Coronavirus ORFeome. PloS One 2007, 2(5), e459. https://doi.org/10.1371/journal.pone.0000459.

(24) de Haan, C. A. M.; Smeets, M.; Vernooij, F.; Vennema, H.; Rottier, P. J. M. Mapping of the Coronavirus Membrane Protein Domains Involved in Interaction with the Spike Protein. J. Virol. 1999, 73(9), 7441–7452.

(25) He, R.; Leeson, A.; Ballantine, M.; Andonov, A.; Baker, L.; Dobie, F.; Li, Y.; Bastien, N.; Feldmann, H.; Strocher, U.; Theriault, S.; Cutts, T.; Cao, J.; Booth, T. F.; Plummer, F. A.; Tyler, S.; Li, X. Characterization of Protein–Protein Interactions between the Nucleocapsid Protein and Membrane Protein of the SARS Coronavirus. Virus Res. 2004, 105(2), 121–125. https://doi.org/10.1016/j.virusres.2004.05.002.

(26) Hsieh, Y.-C.; Li, H.-C.; Chen, S.-C.; Lo, S.-Y. Interactions between M Protein and Other Structural Proteins of Severe, Acute Respiratory Syndrome-Associated Coronavirus. J. Biomed. Sci. 2008, 15(6), 707–717. https://doi.org/10.1007/s11373-008-9278-3.

(27) Kuo, L.; Masters, P. S. Genetic Evidence for a Structural Interaction between the Carboxy Termini of the Membrane and Nucleocapsid Proteins of Mouse Hepatitis Virus. J. Virol. 2002, 76(10), 4987–4999. https://doi.org/10.1128/JVI.76.10.4987-4999.2002.

(28) Kuo, L.; Hurst-Hess, K. R.; Koetzner, C. A.; Masters, P. S. Analyses of Coronavirus Assembly Interactions with Interspecies Membrane and Nucleocapsid Protein Chimeras. J. Virol. 2016, 90(9), 4357–4368. https://doi.org/10.1128/JVI.03212-15.

(29) Fang, X.; Ye, L.; Timani, K. A.; Li, S.; Zen, Y.; Zhao, M.; Zheng, H.; Wu, Z. Peptide Domain Involved in the Interaction between Membrane Protein and Nucleocapsid Protein of SARS-Associated Coronavirus. J. Biochem. Mol. Biol. 2005, 38(4), 381–385. https://doi.org/10.5483/bmbrep.2005.38.4.381.

(30) Hurst, K. R.; Kuo, L.; Koetzner, C. A.; Ye, R.; Hsue, B.; Masters, P. S. A Major Determinant for Membrane Protein Interaction Localizes to the Carboxy-Terminal Domain of the Mouse Coronavirus Nucleocapsid Protein. J. Virol. 2005, 79(21), 13285–13297. https://doi.org/10.1128/JVI.79.21.13285-13297.2005.

(31) Luo, H.; Wu, D.; Shen, C.; Chen, K.; Shen, X.; Jiang, H. Severe Acute Respiratory Syndrome Coronavirus Membrane Protein Interacts with Nucleocapsid Protein Mostly through Their Carboxyl Termini by Electrostatic Attraction. Int. J. Biochem. Cell Biol. 2006, 38(4), 589–599. https://doi.org/10.1016/j.biocel.2005.10.022.

(32) Verma, S.; Bednar, V.; Blount, A.; Hogue, B. G. Identification of Functionally Important Negatively Charged Residues in the Carboxy End of Mouse Hepatitis Coronavirus A59 Nucleocapsid Protein. J. Virol. 2006, 80(9), 4344–4355. https://doi.org/10.1128/JVI.80.9.4344-4355.2006.

(33) Chang, C.; Chen, C.-M. M.; Chiang, M.; Hsu, Y.; Huang, T. Transient Oligomerization of the SARS-CoV N Protein – Implication for Virus Ribonucleoprotein Packaging. PLOS ONE 2013, 8(5), e65045. https://doi.org/10.1371/journal.pone.0065045.

(34) Fernández-Recio, J.; Totrov, M.; Abagyan, R. Identification of Protein–Protein Interaction Sites from Docking Energy Landscapes. J. Mol. Biol. 2004, 335(3), 843–865. https://doi.org/10.1016/j.jmb.2003.10.069.

(35) Ritchie, D. W. Recent Progress and Future Directions in Protein-Protein Docking. Curr. Protein Pept. Sci. 2008, 9(1), 1–15. https://doi.org/10.2174/138920308783565741.

(36) Pan, A. C.; Jacobson, D.; Yatsenko, K.; Sritharan, D.; Weinreich, T. M.; Shaw, D. E. Atomic-Level Characterization of Protein–Protein Association. Proc. Natl. Acad. Sci. 2019, 116(10), 4244–4249. https://doi.org/10.1073/pnas.1815431116.

(37) Walls, A. C.; Park, Y.-J.; Tortorici, M. A.; Wall, A.; McGuire, A. T.; Veesler, D. Structure, Function, and Antigenicity of the SARS-CoV-2 Spike Glycoprotein. Cell 2020, 181(2), 281–292.e6. https://doi.org/10.1016/j.cell.2020.02.058.

(38) Zhang, L.; Lin, D.; Sun, X.; Curth, U.; Drosten, C.; Sauerhering, L.; Becker, S.; Rox, K.; Hilgenfeld, R. Crystal Structure of SARS-CoV-2 Main Protease Provides a Basis for Design of Improved α- Ketoamide Inhibitors. Science 2020, 368(6489), 409–412. https://doi.org/10.1126/science.abb3405.

(39) Hillen, H. S.; Kokic, G.; Farnung, L.; Dienemann, C.; Tegunov, D.; Cramer, P. Structure of Replicating SARS-CoV-2 Polymerase. Nature 2020, 584(7819), 154–156. https://doi.org/10.1038/s41586-020-2368-8.

(40) Yang, J.; Yan, R.; Roy, A.; Xu, D.; Poisson, J.; Zhang, Y. The I-TASSER Suite: Protein Structure and Function Prediction. Nat. Methods 2015, 12(1), 7–8. https://doi.org/10.1038/nmeth.3213.

(41) Kumar, A.; Kumar, P.; Saumya, K. U.; Kapuganti, S. K.; Bhardwaj, T.; Giri, R. Exploring the SARS-CoV-2 Structural Proteins for Multi-Epitope Vaccine Development: An in-Silico Approach. Expert Rev. Vaccines 2020, 0(0), 1–12. https://doi.org/10.1080/14760584.2020.1813576.

(42) Sharma, N.; Prosser, O.; Kumar, P.; Tuplin, A.; Giri, R. Small Molecule Inhibitors Possibly Targeting the Rearrangement of Zika Virus Envelope Protein. Antiviral Res. 2020, 182, 104876. https://doi.org/10.1016/j.antiviral.2020.104876.

(43) Kozakov, D.; Brenke, R.; Comeau, S. R.; Vajda, S. PIPER: An FFT-Based Protein Docking Program with Pairwise Potentials. Proteins 2006, 65(2), 392–406. https://doi.org/10.1002/prot.21117.

(44) Chuang, G.-Y.; Kozakov, D.; Brenke, R.; Comeau, S. R.; Vajda, S. DARS (Decoys As the Reference State) Potentials for Protein-Protein Docking. Biophys. J. 2008, 95(9), 4217–4227. https://doi.org/10.1529/biophysj.108.135814.

(45) Bowers, K. J.; Chow, D. E.; Xu, H.; Dror, R. O.; Eastwood, M. P.; Gregersen, B. A.; Klepeis, J. L.; Kolossvary, I.; Moraes, M. A.; Sacerdoti, F. D.; Salmon, J. K.; Shan, Y.; Shaw, D. E. Scalable Algorithms for Molecular Dynamics Simulations on Commodity Clusters. In SC ‘06: Proceedings of the 2006 ACM/IEEE Conference on Supercomputing; 2006; pp 43–43. https://doi.org/10.1109/SC.2006.54.

(46) Banks, J. L.; Beard, H. S.; Cao, Y.; Cho, A. E.; Damm, W.; Farid, R.; Felts, A. K.; Halgren, T. A.; Mainz, D. T.; Maple, J. R.; Murphy, R.; Philipp, D. M.; Repasky, M. P.; Zhang, L. Y.; Berne, B. J.; Friesner, R. A.; Gallicchio, E.; Levy, R. M. Integrated Modeling Program, Applied Chemical Theory (IMPACT). J. Comput. Chem. 2005, 26(16), 1752–1780. https://doi.org/10.1002/jcc.20292.

(47) Subissi, L.; Posthuma, C. C.; Collet, A.; Zevenhoven-Dobbe, J. C.; Gorbalenya, A. E.; Decroly, E.; Snijder, E. J.; Canard, B.; Imbert, I. One Severe Acute Respiratory Syndrome Coronavirus Protein Complex Integrates Processive RNA Polymerase and Exonuclease Activities. Proc. Natl. Acad. Sci. 2014, 111(37), E3900–E3909. https://doi.org/10.1073/pnas.1323705111.

(48) Kirchdoerfer, R. N.; Ward, A. B. Structure of the SARS-CoV Nsp12 Polymerase Bound to Nsp7 and Nsp8 Co-Factors. Nat. Commun. 2019, 10(1), 1–9. https://doi.org/10.1038/s41467-019-10280-3.

(49) Ma, Y.; Wu, L.; Shaw, N.; Gao, Y.; Wang, J.; Sun, Y.; Lou, Z.; Yan, L.; Zhang, R.; Rao, Z. Structural Basis and Functional Analysis of the SARS Coronavirus Nsp14–Nsp10 Complex. Proc. Natl. Acad. Sci. U. S. A. 2015, 112(30), 9436–9441. https://doi.org/10.1073/pnas.1508686112.

(50) McBride, C. E.; Li, J.; Machamer, C. E. The Cytoplasmic Tail of the Severe Acute Respiratory Syndrome Coronavirus Spike Protein Contains a Novel Endoplasmic Reticulum Retrieval Signal That Binds COPI and Promotes Interaction with Membrane Protein. J. Virol. 2007, 81(5), 2418–2428. https://doi.org/10.1128/JVI.02146-06.

(51) Fung, T. S.; Liu, D. X. Human Coronavirus: Host-Pathogen Interaction. Annu. Rev. Microbiol. 2019, 73(1), 529–557. https://doi.org/10.1146/annurev-micro-020518-115759.

