## Supplementary information for "An insight into SARS-CoV-2 Membrane protein interaction with Spike, Envelope, and Nucleocapsid proteins"

**Supplementary Figures**

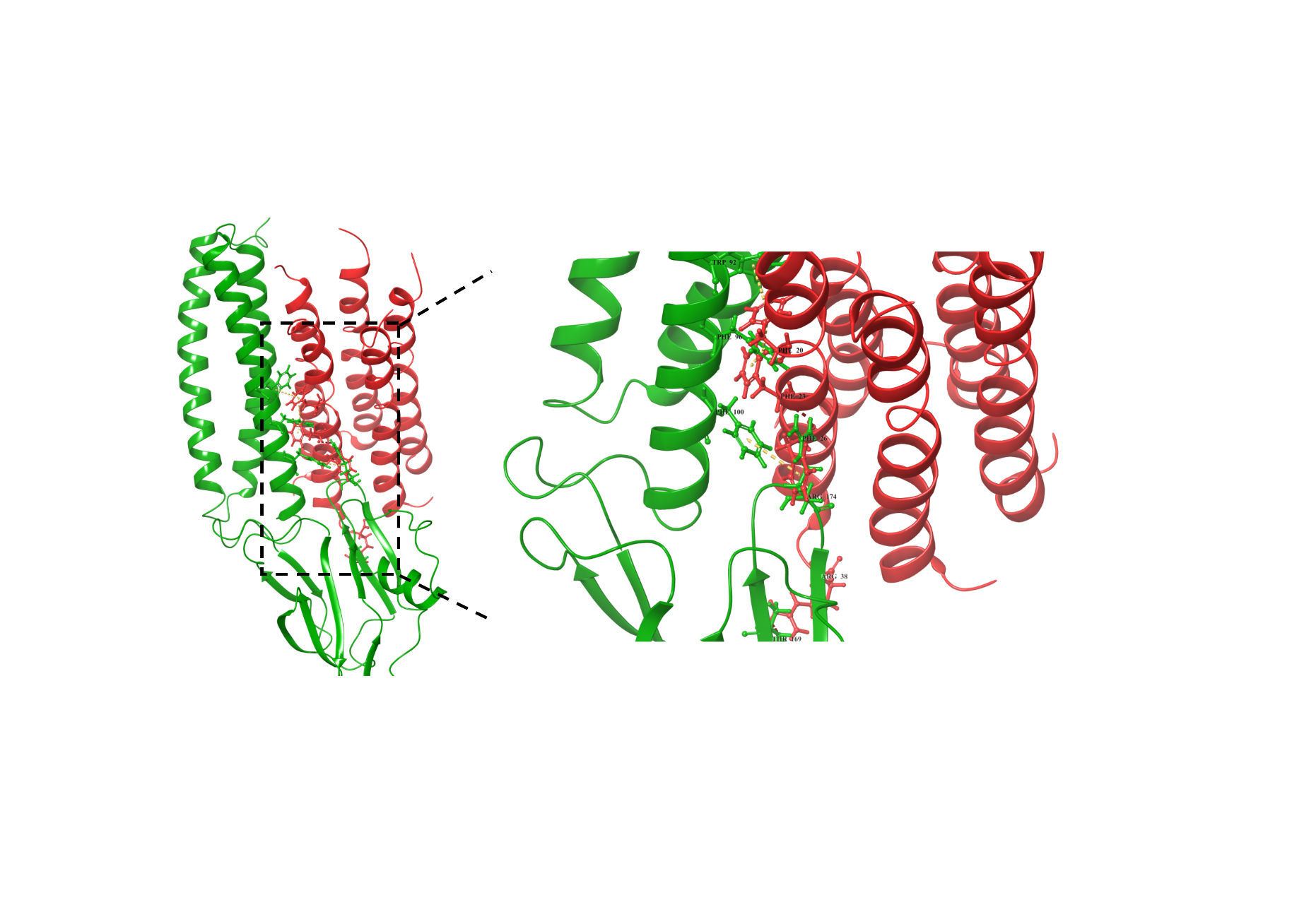

**Supplementary Figure 1:** Molecular interaction analysis of membrane (M) and envelope (E) protein through protein-protein docking. The envelope protein structure is obtained from RCSB-PDB (7K3G) and docked in its pentameric form. Here, the residues Trp92, Phe96, Phe100, Thr169, and Arg174 of M protein are interacting with Phe20, Phe23, Phe26, and Arg38 via 2 hydrogen bonds and 4 π-π stacking contacts.

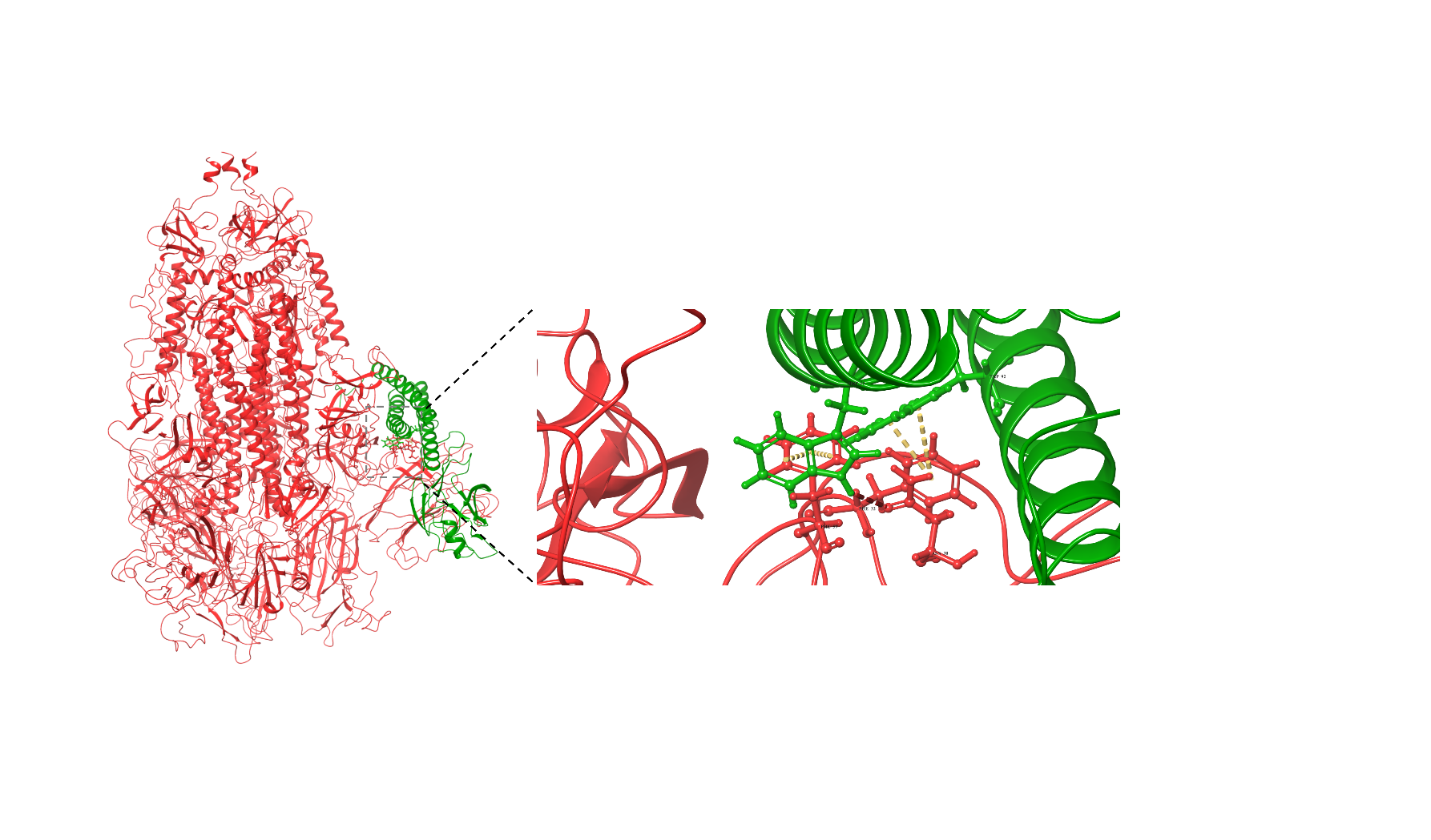

**Supplementary Figure 2:** Molecular interaction analysis of membrane and spike protein through protein-protein docking. The spike protein structure is obtained from RCSB-PDB (6VXX) and docked in its trimeric form. Majorly, M protein interacts through residues through its aromatic residues, Trp55 and Trp92 with E protein (residues Asn30 (hydrogen bond), Phe32 (2x π-π stacking), and Phe59 (2x π-π stacking).

**Supplementary Tables**

**Supplementary Table 1:** Binding energy calculation of protein-protein complexes using MM-GBSA approach. The complexes are selected at every 25ns of simulation trajectory and compared with the docked complex (obtained from protein-protein docking).

| **Protein-protein complex** | **MD Frame** | **Prime energy**  **(kcal/mol)** | **Binding energy**  **(kcal/mol)** |
| --- | --- | --- | --- |
| **Membrane-Envelope** | Docked Complex | -10905.5 | 38.96 |
|  | 25ns | -8399.4 | -273.80 |
|  | 50ns | -8379.8 | -233.79 |
|  | 75ns | -8351.2 | -248.29 |
|  | 100ns | -8330.2 | -261.59 |
| **Membrane-Spike** | Docked Complex | -49369.2 | 10312.30 |
|  | 25ns | -55862.6 | 604.85 |
|  | 50ns | -55999.2 | 713.95 |
|  | 75ns | -55945.8 | 805.55 |
|  | 100ns | -56009.8 | 837.20 |
| **Membrane-Nucleocapsid** | Docked Complex | -25488.1 | -59.80 |
|  | 25ns | -22855.1 | -127.84 |
|  | 50ns | -22859.5 | -55.99 |
|  | 75ns | -22939.2 | -65.74 |
|  | 100ns | -22845.1 | -82.92 |

**Supplementary Table 2:** Protein-protein interaction analysis of frames from 100 ns simulation trajectory of M-E complex.

|  | **Membrane Residue** | **Envelope Residue** | **Interaction Type** |
| --- | --- | --- | --- |
| **25ns** | ALA 2 | TYR 2 | H-bond, Aromatic H-bond |
|  | ASP 3 | MET 1 | H-bond |
|  | ASN 5 | PHE 3 | H-bond |
|  | PHE 45 | SER 55 | Aromatic H-bond |
|  | TRP 55 | PHE 23 | Aromatic H-bond |
|  | PHE 96 | PHE 26 | π-π stacking |
|  | TYR 178 | CYS 40 | H-bond |
|  | SER 214 | CYS 44 | H-bond |
| **50 ns** | MET 1 | ASP 72 | H-bond, Salt bridge |
|  | ALA 2 | ASP 72 | H-bond |
|  | PHE 45 | SER 55 | Aromatic H-bond |
|  | TRP 55 | PHE 23 | H-bond |
|  | TYR 71 | PHE 4 | Aromatic H-bond |
|  | PHE 100 | THR 30 | Aromatic H-bond |
|  | TYR 178 | CYS 40 | Aromatic H-bond |
| **75 ns** | MET 1 | ASP 72 | H-bond, Salt bridge |
|  | ALA 2 | PHE 4 | H-bond |
|  | ALA 2 | ASP 72 | H-bond |
|  | ASP 3 | MET 1 | H-bond |
|  | LEU 51 | PHE 26 | Aromatic H-bond |
|  | TRP 55 | PHE 23 | H-bond |
|  | TRP 55 | PHE26 | π-π stacking |
|  | TRP 92 | PHE 20 | π-π stacking |
|  | TRP 92 | LEU 19 | Aromatic H-bond |
|  | TYR 178 | SER 50 | H-bond |
|  | TYR 178 | CYS 40 | H-bond, Aromatic H-bond |
| **100 ns** | MET 1 | ASP 72 | H-bond, Salt bridge |
|  | ALA 2 | MET 1 | H-bond |
|  | ALA 2 | ASP 72 | H-bond |
|  | PHE 45 | LEU 51 | Aromatic H-bond |
|  | TRP 55 | PHE 23 | H-bond |
|  | ILE 82 | ASN 15 | H-bond |
|  | PHE 96 | PHE 26 | Aromatic H-bond |
|  | PHE 100 | THR 30 | Aromatic H-bond |
|  | TYR 178 | SER 50 | H-bond |
|  | TYR 178 | CYS 40 | H-bond, Aromatic H-bond |
|  | ASN 216 | CYS 43 | H-bond |

**Supplementary Table 3:** Protein-protein interaction analysis of frames from 100 ns simulation trajectory of M-S complex.

|  | **Membrane Residue** | **Spike Residue** | **Interaction Type** |
| --- | --- | --- | --- |
| **25 ns** | ASN 5 | ILE 197 | H-bond |
|  | LYS 15 | CYS 166 | Salt bridge |
|  | GLN 19 | ASN 234 | H-bond |
|  | CYS 86 | LEU 10 | H-bond |
|  | PHE 100 | THR 323 | Aromatic H-bond |
|  | ARG 101 | GLU 324 | H-bond (2), Salt bridge |
|  | ARG 174 | CYS 538 | Salt bridge |
|  | ARG 174 | GLN 321 | H-bond |
|  | ARG 174 | GLN 628 | H-bond |
|  | ASN 203 | THR 618 | H-bond |
|  | ASN 207 | ASN 616 | H-bond |
| **50 ns** | ASN 5 | ASP 198 | H-bond |
|  | THR 7 | THR 167 | H-bond |
|  | THR 7 | TYR 200 | Aromatic H-bond |
|  | LYS 15 | CYS 166 | Salt bridge |
|  | GLN 19 | ASN 234 | H-bond |
|  | CYS 86 | LEU 10 | H-bond |
|  | PHE 96 | VAL 16 | Aromatic H-bond |
|  | PHE 100 | THR 323 | Aromatic H-bond |
|  | ARG 101 | GLU 324 | H-bond (2), Salt bridge |
|  | ARG 174 | CYS 538 | Salt bridge |
|  | ARG 174 | GLN 321 | H-bond |
|  | ARG 174 | GLN 628 | H-bond |
|  | ASN 203 | THR 618 | Aromatic H-bond |
|  | ASN 207 | ASN 616 | Aromatic H-bond |
| **75 ns** | ASN 5 | ASP 198 | H-bond |
|  | THR 7 | THR 167 | H-bond |
|  | THR 7 | TYR 200 | H-bond, Aromatic H-bond |
|  | LEU 16 | ASN 234 | H-bond |
|  | CYS 86 | LEU 10 | H-bond |
|  | PHE 96 | VAL 16 | Aromatic H-bond |
|  | PHE 100 | THR 323 | Aromatic H-bond |
|  | ARG 101 | GLU 324 | H-bond, Salt bridge |
|  | ARG 174 | CYS 538 | Salt bridge |
|  | ARG 174 | GLN 321 | H-bond |
|  | ARG 174 | GLN 628 | H-bond (2) |
|  | LYS 205 | CYS 617 | H-bond |
|  | ASN 207 | ASN 616 | H-bond |
| **100ns** | SER 4 | ASP 198 | H-bond |
|  | ASN 5 | ASP 198 | H-bond |
|  | LEU 16 | ASN 234 | H-bond |
|  | TRP 20 | ASN 87 | Aromatic H-bond |
|  | PH 96 | VAL 16 | Aromatic H-bond |
|  | SER 173 | LYS 537 | H-bond |
|  | SER 173 | THR 323 | H-bond |
|  | ARG 174 | CYS 538 | H-bond, Salt bridge |
|  | ARG 200 | GLU 554 | H-bond (2), Salt bridge |

**Supplementary Table 4:** Protein-protein interaction analysis of frames from 100 ns simulation trajectory of M-N complex.

|  | **Membrane Residue** | **Nucleocapsid Residue** | **Interaction Type** |
| --- | --- | --- | --- |
| **25 ns** | ARG 150 | GLU 323 | H-bond |
|  | ARG 150 | VAL 324 | H-bond |
|  | ASN 207 | PHE 286 | Aromatic H-bond |
|  | HIS 210 | GLN 272 | H-bond |
|  | HIS 210 | ALA 313 | Aromatic H-bond |
|  | HIS 210 | THR 271 | Aromatic H-bond |
|  | ASP 215 | ALA 336 | H-bond |
| **50 ns** | ARG 150 | GLU 323 | H-bond |
|  | ARG 150 | VAL 324 | H-bond |
|  | SER 197 | THR 334 | H-bond |
|  | THR 208 | PHE 314 | Aromatic H-bond |
|  | ASP 215 | ALA 336 | H-bond |
| **75 ns** | ASN 207 | MET 317 | H-bond |
|  | THR 208 | TYR 333 | Aromatic H-bond |
|  | THR 208 | PHE 314 | Aromatic H-bond |
|  | HIS 210 | TYR 333 | H-bond |
|  | SER 211 | THR 334 | H-bond |
|  | ASP 215 | ALA 336 | H-bond |
| **100 ns** | ARG 150 | GLU 323 | H-bond, Salt bridge |
|  | GLY 192 | PRO 326 | H-bond |
|  | HIS 210 | GLN 272 | Aromatic H-bond |
|  | THR 208 | PHE 314 | Aromatic H-bond |
|  | ASN 207 | PHE 286 | Aromatic H-bond |
|  | PHE 193 | TRP 330 | π-π stacking |

**Supplementary Movies**

Movie 1: Simulation trajectory movie of M-E docked complex upto 100 ns timescale.

Movie 2: Simulation trajectory movie of M-S docked complex upto 100 ns timescale.

Movie 3: Simulation trajectory movie of M-N docked complex upto 100 ns timescale.
